# Effect of an amyloidogenic SARS-COV-2 protein fragment on α-synuclein monomers and fibrils

**DOI:** 10.1101/2022.02.21.481360

**Authors:** Asis K. Jana, Chance W. Lander, Andrew D. Chesney, Ulrich H. E. Hansmann

## Abstract

Using molecular dynamic simulations we study whether amyloidogenic regions in viral proteins can initiate and modulate formation of α-synuclein aggregates, thought to be the disease-causing agent in Parkinson’s Disease. As an example we choose the nine-residue fragment SFYVYSRVK (SK9), located on the C-terminal of the Envelope protein of SARS-COV-2. We probe how the presence of SK9 affects the conformational ensemble of α-synuclein monomers and the stability of two resolved fibril polymorphs. We find that the viral protein fragment SK9 may alter α-synuclein amyloid formation by shifting the ensemble toward aggregation-prone and preferentially rod-like fibril seeding conformations. However, SK9 has only little effect of the stability of pre-existing or newly-formed fibrils.

## Introduction

While most COVID-19 survivors seem to recover fully, not much is known about potential long-term or delayed neurological effects of SARS-COV-2 infections. Acute infection-associated symptoms include loss of smell and other neurological impairments^1^; and a number of case studies suggests a role of SARS-CoV-2 in the onset of Parkinson’s Disease and related neurodegenerative diseases^2–5^. A possible mechanism for triggering Parkinson’s Disease could be virus-initiated amyloid-formation as aggregates of α-synuclein (aS) are the cell-toxic agent in Parkinson’s Disease and related pathologies^6^. SARS-COV-2 accelerated aS amyloid formation, caused by interaction with amyloidogenic regions on the Spike, Envelope or Nucleocapsid protein, has been observed *in vitro*^4^, and may not be accidental. It has been speculated in the context of Alzheimer’s Disease that amyloid fibrils form as an immune response to an infection, entrapping and neutralizing pathogens^7–9^; and a similar mechanism may also play a role in Parkinson’s Disease. However, the link between exposure to pathogens, appearance of fibrils, and subsequently of disease symptoms is, even in the case of Alzheimer’s Disease, only indirect. In order to evaluate whether such a microbial protection hypothesis^7,8^ can explain the correlations between COVID-19 and outbreak of Parkinson’s Disease, one needs to understand the mechanism by which amyloidogenic regions on SARS-COV-2 proteins initiate and modulate aggregation of α-synuclein.

In the present paper we focus on the nine-residue fragment SFYVYSRVK (SK9) on the C-terminal of the Envelope protein. We are familiar with SK9 from other work,^10,11^ and study now its effect on α-synuclein amyloid formation. The 140-residue long α-synuclein monomer consists of three domains: (i) an amphipathic N-terminal region (residues 1-60), where most disease-causing familial mutations are located and which binds to membranes; (ii) a hydrophobic non-amyloid-β component (NAC) region (residues 61-95) that is thought to be the core region for aggregation; and (iii) a disordered negatively charged C-terminal region (residues 96-140).^12–15^ A helix-rich model of the α-synuclein monomer structure has been resolved by solution NMR in the micellar environment and is deposited in the Protein Data Bank (PDB) under PDB ID: 1XQ8.^12^ Using molecular dynamics simulations we study how SK9 interacts with α-synuclein and whether SK9 changes the conformational ensemble of the monomer, potentially modifying oligomerization and hastening the formation of a critical nucleus for seeding fibrils.

In a complementary set of simulations we explore whether and how SK9 differentially alters the stability of experimentally resolved α-synuclein fibril models, comparing the two predominant forms, the “rod” or “twister” polymorphs.^16^ While in both structures the individual chains share a bent β-arch architecture, they have different inter-protofilament interfaces. In the twister polymorph is the interface formed by the hydrophobic aggregation-inducing non-amyloid-β component (NAC) region (G68-A78), while in the rod polymorph the interface is made of the preNAC region(E46-A56).^16,17^ The residues at the C-terminus in the rod polymorph are more ordered, suggesting higher stability than found for the twister polymorph.^16^ Note, however, that the six common mutations E46K, H50Q, G51D, A53E, A53T, and A53V, destabilize the preNAC interface in the rod structure while not disrupting the twister polymorph, i.e., likely shifting the population from rod to twister.^16,18^ As these mutations are associated with hereditary forms of Parkinson’s Disease, it appears as if a shift in frequency between rod and twister fibril forms will alter the probability for acquiring Parkinson’s Disease.^16,18^

Our goal in the present paper is therefore to study if and how the viral segment SK9 accelerates aggregation of α-synuclein, and whether this process is selective, preferentially enhancing the formation of potentially more toxic amyloid species.

## Materials and Methods

### System Preparation

In order to understand the effect of SARS-COV-2 proteins on α-synuclein aggregation we use molecular dynamics simulations to study how the presence of the nine-residue segment S^55^FYVYSRVK^63^ (SK9) of the Envelope protein changes the conformational ensemble and stability of α-synuclein monomers and fibrils. For the initial configuration for the monomer, we chose the model resolved by solution NMR and deposited in the Protein Data Bank (PDB) under PDB ID: 1XQ8. As this structure is characterized by a largely helical N-terminus,^12^ and to avoid any bias by this choice of an initial configuration, we have also considered a fully extended α-synuclein monomer as an alternative start configuration. It is obtained by heating the experimentally resolved conformation up to 500 K, and simulating it for 5 ns. On the other hand, for our investigations into the stability of α-synuclein fibrils, we have considered the two predominant forms, the rod and a twister polymorphs. For this purpose, we have generated decamers, made of two folds (protofilaments) and five layers, from the cryo-EM structures deposited in the PDB under identifier 6CU7 and 6CU8, respectively.^16^ Note that for the rod model only 60 residues (L38-K97) out of the 140 amino acids in α-synuclein have been resolved, and only 41 residues (K43-E83) for the twister model. For this reason, we have also generated models for the rod and the twister fibril where all of the experimentally unsolved regions are added as predicted by homology modeling. However, since subsequent docking experiments did not indicate binding of the SK9 segment to the N-terminal residues 1-37 or the C-terminal residues 121-140, we decided to reduce computational costs by only considering extended rod and twister fibril models made of residues 38-120. For both monomer and fibril models, the N- and C-terminus of the chains are capped by NH3^+^ and COO^-^ groups, respectively.

Simulations starting from the two monomer and four fibril models serve as controls against simulations in which SK9 is present. The initial configuration for the SK9 fragment was derived by removing the residues 1-54 and 64-75 from the model of the SARS-COV-2 E-protein as deposited in the MOLSSI COVID-19 hub^19^, with the N- and C-termini of the peptide capped by a NH3+ and -CONH2 group, respectively. Using the AutoDock Vina software^20^, we have generated start configurations for our simulations by docking SK9 segments in a ratio of 1:1 with the α-synuclein chains in the monomer and fibril systems described above. The so-generated six structures are shown in **Figure 1**, but note that SK9 segment is not constrained to these positions and can move freely (and even detach) in the simulations.

**Figure 1:**
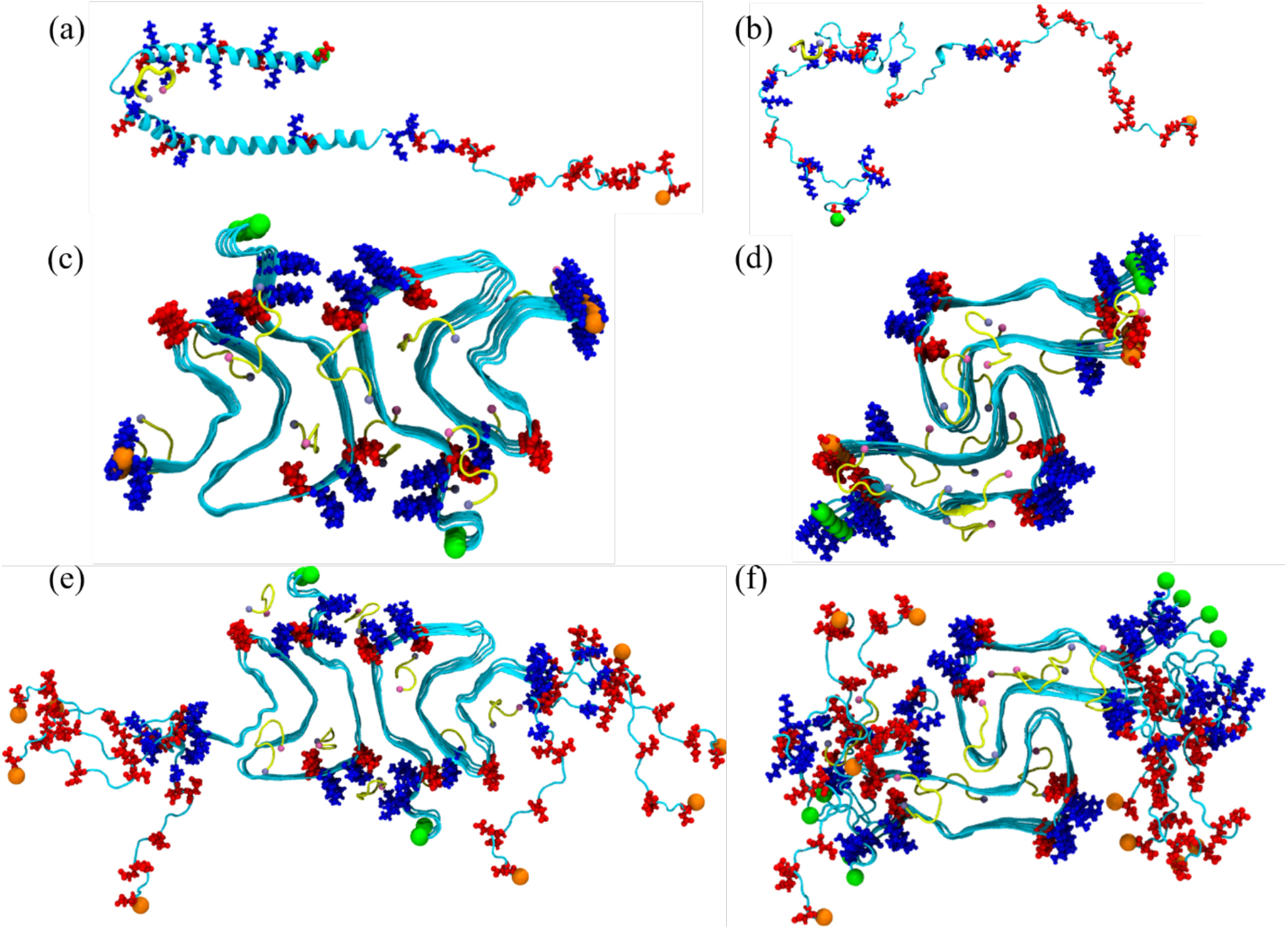
Initial conformation of the α-synuclein monomer (a) as resolved by solution NMR (PDB ID: 1XQ8), and (b) after heating at 500 K to obtain a randomized stretched conformation. Initial conformation for the fibril as derived by cryo-EM structures are shown in (c) for the rod (PDB ID: 6CU7) and in (d) for the twister (PDB ID: 6CU8) polymorph. In (e) and (f) are the corresponding structures shown for the fibrils where the individual chains are extended to residues 38-120. Acidic residues are colored in red and basic ones in blue, while the SK9-segments are shown yellow. The N- and C-termini are represented by green and orange spheres, respectively.

The twelve models of α-synuclein monomers and fibrils, either with SK9 binding to the α-synuclein chain or with SK9 absent, are each put in a rectangular box with periodic boundary conditions. The box is chosen such that there is a minimum distance of 15 Å between any SAA or SK9 atom and all box sides. This large box size ensures that the α-synuclein chains do not interact with their periodic images. Each box is filled with water molecules, and counterions are added to neutralize the system. Additional Na+ and Cl− ions were added to arrive at a physiological ion concentration of 0.15 M NaCl. **Table 1** lists for all systems the number of water molecules and the total number of atoms.

**Table 1:** Simulated systems.

| system description |  | atoms | water molecules | Independent trajectories | simulation length | total sampling |
| --- | --- | --- | --- | --- | --- | --- |
| α-synuclein1-140 monomer, (PDB-ID:1XQ8 model) | w/o SK9 | 287,045 | 71,155 | 3 | 3.0 μs | 9.0 μs |
| | w SK9 | 288,809 | 71,508 | 3 | 3.0 μs | 9.0 μs |
| α-synuclein1-140 monomer, (fully extended conformation) | w/o SK9 | 284,469 | 70,512 | 2 | 3.0 μs | 6.0 μs |
| | w SK9 | 329,376 | 81,683 | 2 | 3.0 μs | 6.0 μs |
| Rod fibril model (PDB ID: 6CU7, resolved for residues 38-97) | w/o SK9 | 183,222 | 58,098 | 3 | 0.2 μs | 0.6 μs |
| | w SK9 | 183,383 | 57,585 | 3 | 0.2 μs | 0.6 μs |
| Extended rod fibril (residue 38-120) | w/o SK9 | 516,415 | 168,653 | 3 | 0.2 μs | 0.6 μs |
| | w SK9 | 516,587 | 168,273 | 3 | 0.2 μs | 0.6 μs |
| Twister fibril model (PDB ID: 6CU8, resolved for residues 43-83) | w/o SK9 | 177,275 | 56,889 | 3 | 0.2 μs | 0.6 μs |
| | w SK9 | 177,583 | 56,525 | 3 | 0.2 μs | 0.6 μs |
| Extended twister fibril (residue 38-120) | w/o SK9 | 324,814 | 104,900 | 3 | 0.2 μs | 0.6 μs |
| | w SK9 | 325,049 | 104,541 | 3 | 0.2 μs | 0.6 μs |
| All trajectories | | | | | | 34.8 μs |

### General Simulation Protocol

All simulations were carried out using the GROMACS 2018 simulation package.^21^ Intramolecular interactions are described for the fibril by the CHARMM 36m all-atom force field^22^ and the TIP3P water model^23^, which we found in earlier work well-suited for simulations of fibril and oligomers.^24,25^ However, the monomer simulations rely on the a99SB-disp all-atom force field and the corresponding a99SB-disp water model. This is because CHARMM and AMBER force fields have been shown to lead to conformational ensembles of α-synuclein chains that are too compact^26^, while the a99SB-disp all-atom force field and a99SB-disp water model lead to ensembles that do agree with the experimental measurements^27^. Prior to the production run, each system is first energy minimized for up to 5 000 000 steps using steepest descent, and equilibrated by 200 ps molecular dynamics in a NVT ensemble of 310 K, and 200 ps in a NPT ensemble at 310 K and 1 atm, with the heavy atoms of α-synuclein chains and SK9 chains constraint with a force constant of 1000 kJ mol^−1^ nm^−2^.

Production simulations are started from the above generated initial conformations and run at 310 K and 1 atm. Temperature is controlled by a v-rescale thermostat with a coupling constant of 0.1 ps, and pressure kept constant by using the Parrinello-Rahman barostat with a pressure relaxation time of 2 ps. Water molecules are kept rigid using the SETTLE algorithm^28^, and non-water bonds involving hydrogen atoms are restrained to their equilibrium lengths using the LINCS^29^ algorithm, allowing us to use a time step of 2 fs for integrating the equations of motions. Long-range electrostatic interactions are, because of the periodic boundary conditions, computed with the particle-mesh Ewald (PME) technique, using a real-space cutoff of 12 Å and a Fourier grid spacing of 1.6 Å. Short-range van der Waal interactions are truncated at 12 Å, with smoothing starting at 10.5 Å. For each setup, we follow three independent trajectories starting from different initial velocity distributions. The position of the SK9 segment is not restrained to the original docking sites. Instead, it can move and is even allowed to detach over the course of the simulations. The length of the various trajectories is also listed in **Table 1**.

### Trajectory Analysis

For most of our analyses we rely on the GROMACS tools and VMD software. For calculating root mean square deviation (RMSD) and root mean square fluctuation (RMSF) with respect to the experimentally solved structure we use the GROMACS tools gmx_rms and gmx_rmsf, respectively. The radius of gyration is calculated using the gmx_gyrate utility, while the residue-wise secondary structural propensity is computed with the Dictionary of Secondary Structure in Proteins (DSSP) as implemented in the GROMACS do_dssp tool. The solvent accessible surface area (SASA) and residue-residue contact maps are computed with the VMD software, using a spherical probe of 1.4 Å radius for SASA calculations, and defining contacts by a cutoff of 7.0 Å in the closest distance between heavy atoms in a residue pair.

## Results

### 1) MONOMER SIMULATIONS

The purpose of this paper is to explore whether and how SARS-COV-2 proteins can change the amyloid formation of α-synuclein. We first look into the effect of the SK9 segment of the Envelope protein on early steps in the oligomerization of α-synuclein. Comparing molecular dynamics simulations of the protein in the presence of SK9 with such where SK9 is absent (our control), we study how the ensemble of α-synuclein monomer conformations is altered by presence of SK9, incrasing or decreasing the likelihood of amyloid formation. The trustworthiness of our results depends on a faithful description of the α-synuclein monomer (i.e., sufficiently accurate force fields), and sufficient sampling of the various systems. Hence, in order to ensure that our data are reliable and not biased by our set-up, we first compare our simulations of the control with experimental data. While α-synuclein is an intrinsically disordered protein, some structural information has been obtained from NMR experiments^30^. These chemical shift data can be compared with such generated from our simulations using the ShiftX software^31^. We show in **Figure 2a** both the experimentally measured values and the ones calculated from our simulations. The corresponding percentage changes for each residue are shown in **Figure 2b**, where the black line marks zero change. While these figures show a systematic deviation, the percentual change is small and the regression plots in the **Supplemental Figure S1** lead to large Pearson correlation coefficients. We remark that the deviations between numerical and NMR measurements are consistent with previously reported coarse-grained molecular dynamic studies^32,33^, and may arise from either systematic errors in the NMR measurements due to reference quantification or a dilution effect^34^, or from use of non-polarizable force fields, leading to systematic errors in the calculations due to the absence of polarization in the H_α_ atoms^35^. Note that, the chemical shift values do not differ between the various trajectories of the isolated α-synuclein monomer suggesting sufficient sampling to achieve equilibrium.

**Figure 2:**
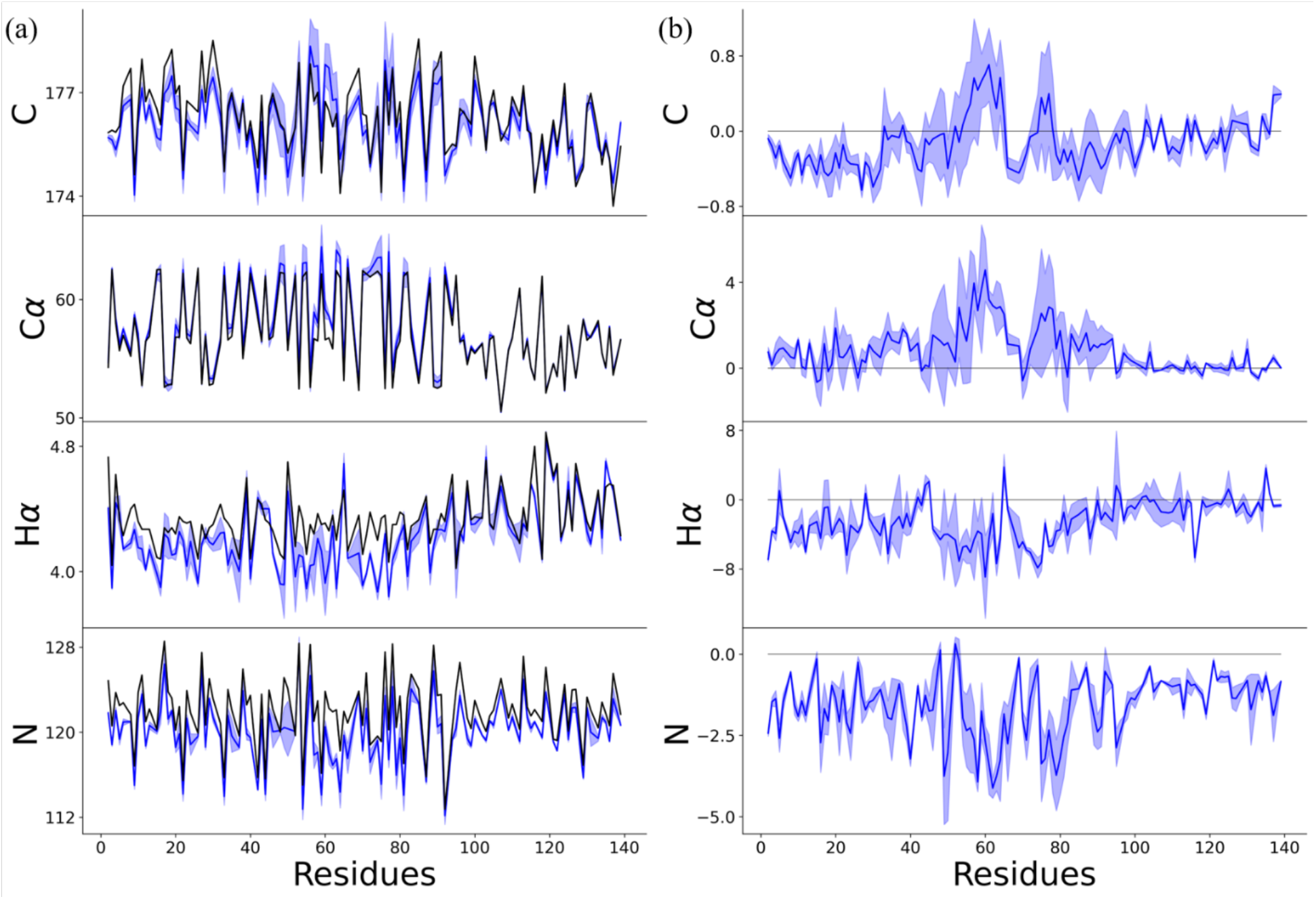
(a) Chemical shifts of carbonyl carbons (C), α-carbons (Cα), hydrogen atoms attached to α-carbons (Hα), and amide nitrogens (N) for α-synuclein monomer, as taken from NMR^30^ (in black) and simulations (in blue). Percentage change in chemical shift values with respect to experimentally measured values are shown in (b) with the black line marking zero deviation.

Our above analysis shows that our simulation set-up describes faithfully the ensemble of α-synuclein monomers and gives us confidence in the reliability of the data measured in these simulations. First, we look into the overall behavior of the chains. Visual inspections indicates that in the presence of SK9, the α-synuclein monomers are more extended and strand-like. This observation is supported by **Figure 3** where we show in **Figure 3a** the distribution of the radius of gyration (Rg), in **Figure 3b** the distribution of the solvent accessible surface area (SASA) and in **Figure 3c** that of the number of contacts (n_C_). We observe in all cases the same trend: the ensemble is in the presence of SK9 shifted toward larger, more solvent-exposed and looser-packed conformations. For the solvent accessible surface area, the effect does not result from increased exposure of hydrophilic residues as it is also seen when the calculation is restricted to hydrophobic residues (in which case the SASA value changes from (4404 ± 522) Å^2^ in the control to (4651 ± 494) Å^2^). Hence, the presence of SK9 shifts the distribution of α-synuclein conformations toward an ensemble with conformations that are more extended than seen in the control, and that expose more hydrophobic residues, making the conformations more aggregation prone.

**Figure 3:**
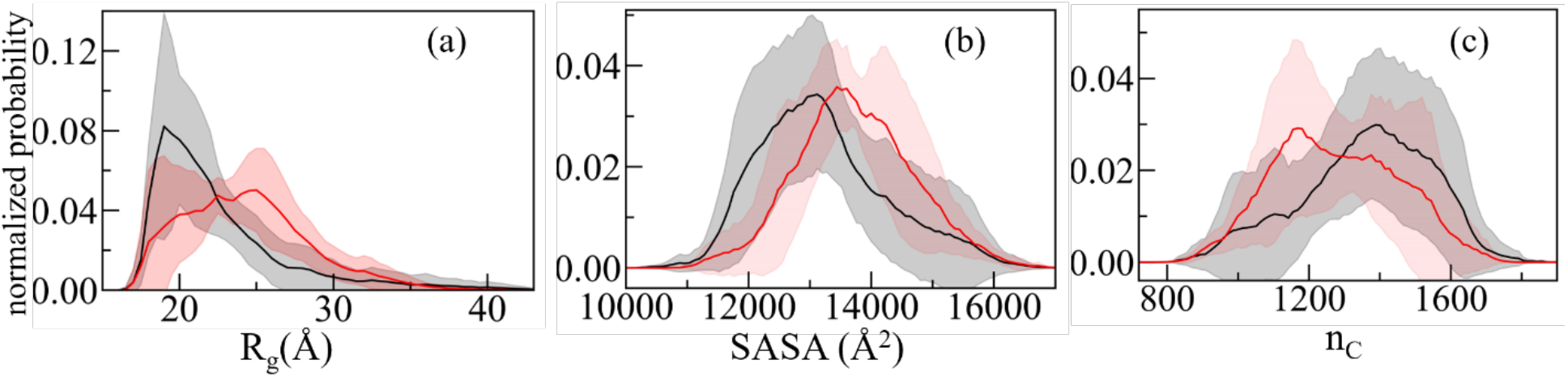
Normalized distribution of (a) radius of gyration (R_g_), (b) solvent accessible surface area (SASA) and (c) the number of contacts (n_C_) of α-synuclein monomer in absence (black) and presence (red) of the SK9-segment. Data are averaged over the final 2.0 *µ*s of each trajectory.

In order to understand how interaction with the viral protein fragment SK9 moves the ensemble of α-synuclein monomer conformations toward more aggregation prone conformations, we show in **Figure 4a** the root-mean-square-fluctuations (RMSF) of the residues, a quantity that measures the flexibility of a given residue. Comparing values measured in simulations that include SK9 with simulations where SK9 is absent, allow us therefore to identify the regions in the α-synuclein chain affected by SK9. **Figure 4a** shows that the α-synuclein monomer is more flexible in the simulations where the viral protein segment is present. An exception is the C-terminal region (residues 100 – 140), which is disorganized in the solution NMR structure of 1XQ8 and which corresponds to the main binding site of SK9, see **Figure 4b**. However, in the other two segments that have enhanced binding probability, residues Q62 to G68 and residues T92 to V95, both located in the NAC region (residues E61 to V95), the binding leads to a higher flexibility and to reduced helicity, see Supplemental **Figure 2a**. As a consequence, the average helicity in the NAC region is reduced from approximately 13% to about 7%. Only a small reduction in helicity (from 5% to 4%) is seen for residues A27-T54, where most of the known familial mutations are located and which is assumed to play a key-role in the early stages of α-synuclein mis-folding and aggregation.^32,36–38^ Similar to the NAC region of residues E61-V95 this region is also characterized by an increased tendency for β-hairpin formation. In the absence of the viral segment SK9, the β-hairpin is transient and is observed with a frequency of less than 11% for residues A27-T54 and for the NAC region. On the other hand, in the presence of SK9, the β-hairpin is formed at an earlier stage and is found in the two regions with a frequency of 15% and 14%, respectively.

**Figure 4:**
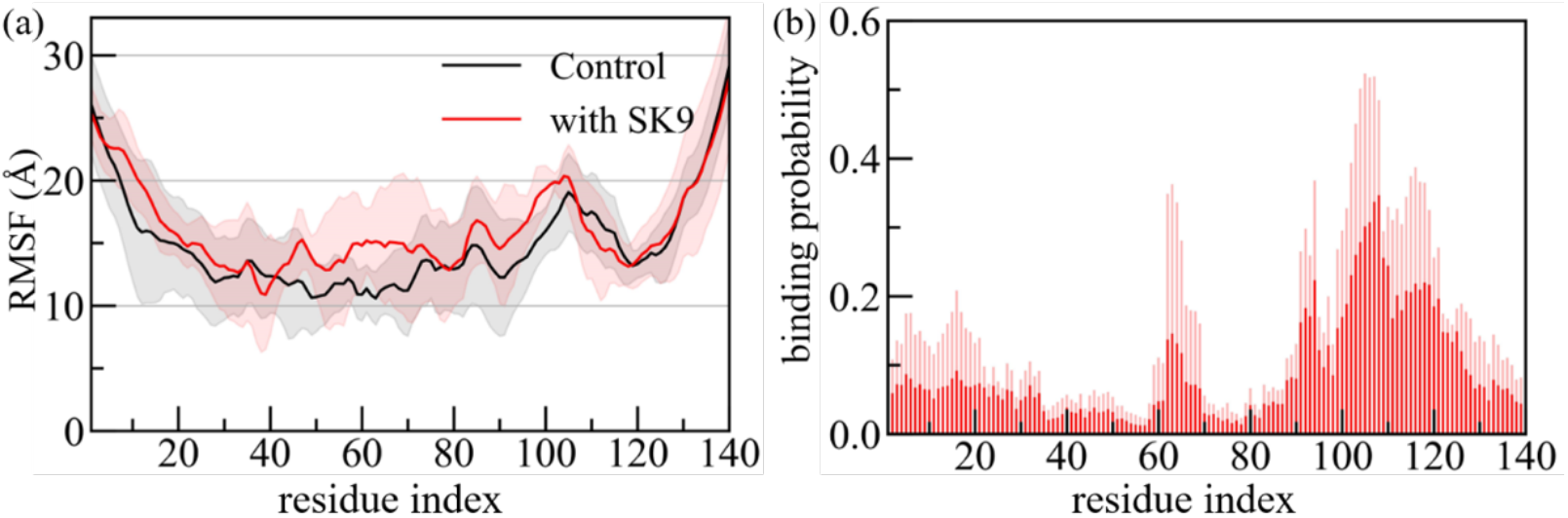
(a) Residue-wise mean square fluctuation (RMSF; in Å) obtained from α-synuclein monomer simulations in the absence (in black) and in the presence (in red) of the SK9-segment. The RMSF values are calculated with reference to the experimentally resolved α-synuclein monomer (PDB ID: 1XQ8) considering only backbone atoms. The residue-wise normalized binding probability of the SK9 segment for α-synuclein monomer is shown in (b). Data are averaged over the final 2.00 μs of each trajectory. Shaded region mark the standard deviation of the averages.

The changes in secondary structure and in the biophysical properties of the α-synuclein monomer results from the interaction of the SK9 segment with the above discussed residues. These interactions modify the network of contacts that stabilize the monomer. Hence, we show in **Figure 5a** and **5b** the residue-residue plot of the contact frequencies observed in our simulations for the control, and in presence of SK9. We show in **Figure 5c** the difference of frequencies of contacts between the two cases, and in **Figure 5d** the difference in their life-times. Note that we show only data for contacts appearing with a frequency of at least 50%. In the absence of SK9 most of the contacts are formed in the familial mutation sites and NAC region, with only low probability to form contacts at the N-terminus and disordered C-terminus. Upon SK9-binding, α-synuclein loses the contacts between residues T44-T54 and A69-V77, located at the familial mutation sites and in the central part of the NAC region, with the average life time of contacts between these regions reduced by about 1.7 ns. A similar loss in contacts is observed between residues K45-K60 and residues K80-G93, where the average lifetime of such contacts is also lowered by about 2.4 ns. Loss of these contacts and such between residues E61-T75 and residues K80-V95 increases the solvent exposure of these regions and also reduces α-helix propensity of the familial mutation sites and in the NAC region while increasing their flexibility, as also seen in **Figure 3** and **Supplemental Figure S2**. On the other hand, the presence of SK9 increases charged and polar interaction within the NAC region involving the residue pairs K60-Q79, E61-K80, Q62-Q79, T64-T75, N65-Q79 and to a smaller extent hydrophobic interactions involving the residue pairs V63-V77, V66-V74 and V66-V77. Together with newly formed contacts between the residues in the segment T33-S42 and with a subset of the NAC region (T33-T75, V37-V77, V37-V82, L38-V82, Y39-T81, V40-V82 and S42-T81) these new interactions cause the structural rearrangement of α-synuclein monomer with its higher β-hairpin frequency and resulting increased probability for aggregation.

**Figure 5:**
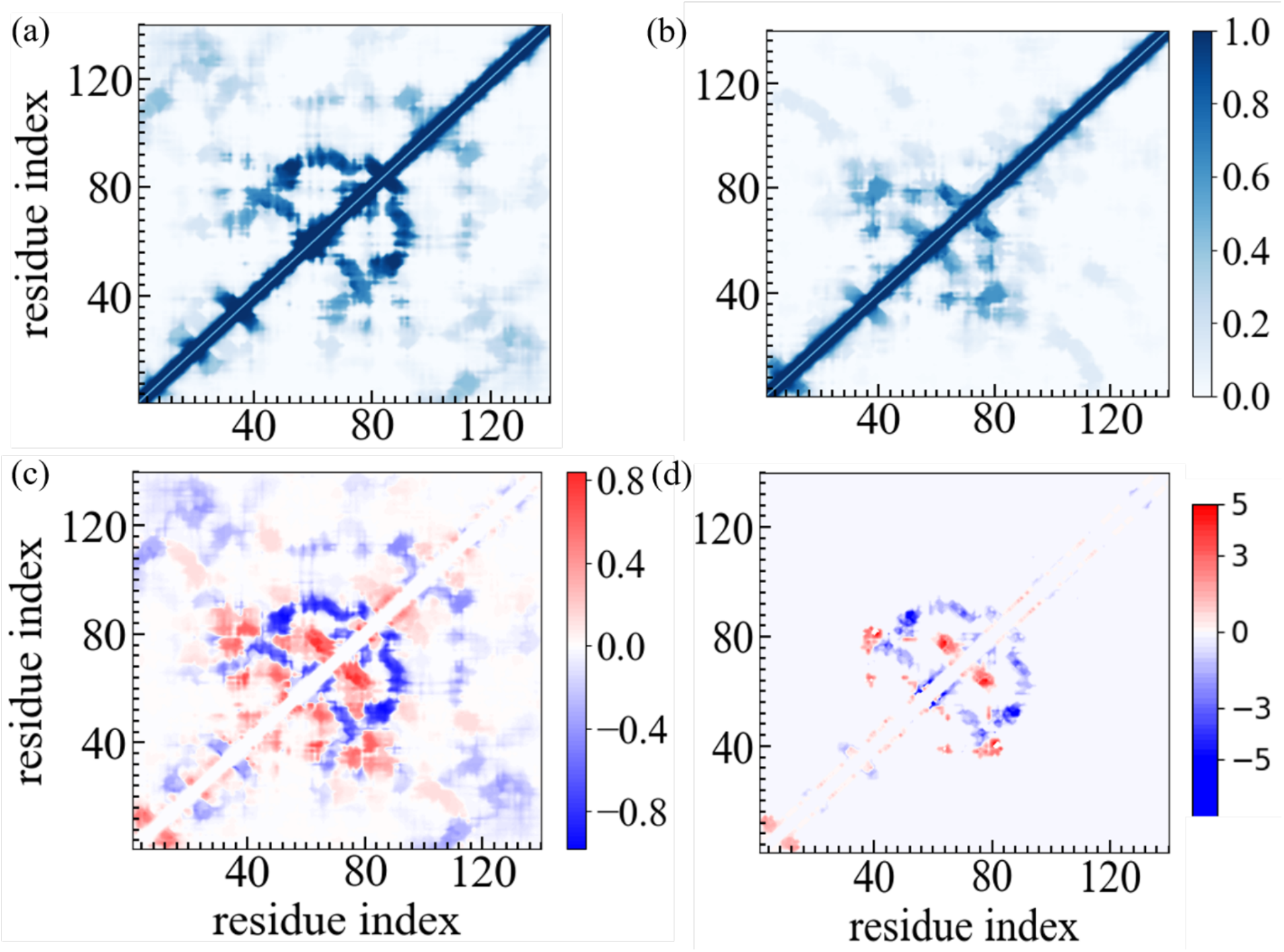
Residue–residue contact frequencies of α-synuclein monomer in (a) the absence and (b) in the presence of the SK9 segment. Data were averaged over the final 2.0 μs of each trajectory. The differences between the two frequencies are shown in (c) while we plot in (d) the corresponding difference in the average lifetime of contacts, i.e, how much longer on average a contact persists in presence of SK9 compared to its average lifetime in the control simulation.

Note that in presence of SK9 the residues in the monomer that form in the rod fibril polymorph (see **Figure 6**) the inter-protofilament interface (E46-A56) are exposed, thereby increasing the chance to bind with another α-synuclein chain in this region. This suggests that the presence of SK9 raises preferentially the likelihood for forming the rod polymorph. This assumption is supported by a comparison of the frequency of contacts that are also observed in either the twister or the rod fibril motif. Two contacts, T54-T75 and E61-T72 are found in both fibril polymorphs, and are seen in the monomer simulations with 7% frequency, with the frequency not affected by the presence of SK9. SK9 also does not noticeably change in the monomer simulations the frequency of two contacts found solely in the rod polymorph: A69-A91 and V77-I88. Both are seen in about 40% of configurations. On the other hand, the presence of SK9 strongly increases the frequency, and doubles the lifetime, of two other stabilizing contacts that are seen in the rod fibril: E46-K80 (from 5% to 36%) and V52-A76 (from 10% to 50%). These two contacts connect residues located in the NAC region thereby fixating the inter-protofilament interface in the rod structure. While we see a similar increase in frequency of contacts in the presence of SK9, from about 16% to 32% between the residue pairs V49-Q79 and V49-V77, which stabilize the inter-protofilament interface in the individual chains in the twister, there is no noticeable change in their average lifetime of about 1 ns. However, the increase in frequency of the two stabilizing contacts is partially compensated by a lower frequency of the contact E46-T81 which, competing with the E46-K80 contact, decreases from 27% to 6%, and a decrease in the frequency of the V63-V70 contact from 57% in the control to 23%. While the average lifetime of the E46-T81 contact decreases from 1.5 ns to 0.3 ns, it stays unchanged at around 1 ns for the V63-V70 contact. Hence, our analysis of the monomer simulations suggests not only that the presence of the SARS-COV-2 protein fragment SK9 raises the probability for α-synuclein amyloid formation, but also that this effect favors the rod motif.

**Figure 6:**
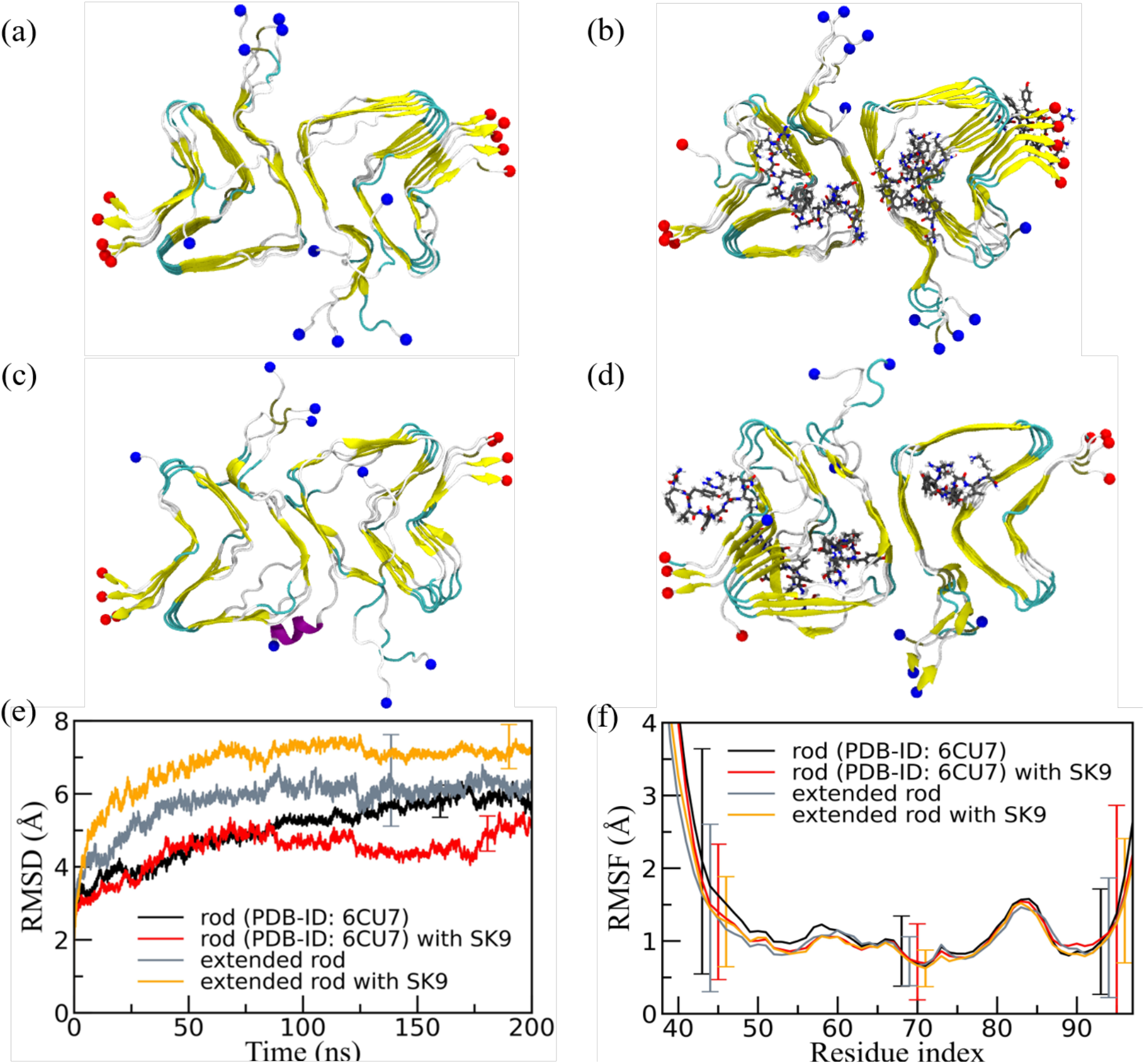
Representative final configurations extracted from simulations starting from (a) the experimentally determined rod-like α-synuclein fibril model (PDB-ID: 6CU7) and (c) the extended model. Corresponding final snapshots extracted from simulations in the presence of SK9-segment are shown in (b), and (d). N- and C-terminus are represented by blue and red spheres, respectively. Only residues 38-97 are shown for the extended model configurations in (c) and(d). The time evolution of the RMSD in the simulation of these systems is shown in (e), and residue-wise RMSF in (f). We calculate RMSD and RMSF again only for the experimentally resolved region 38-97, i.e., ignoring the disordered and unresolved parts of the fibril models, considering all backbone atoms. Only a few typical errorbars are shown to make figures more readable.

### 2) FIBRIL SIMULATIONS

The net-effect of the interaction between SK9 and α-synuclein monomers is a shift in the conformational ensemble toward monomer conformations that are more prone to aggregate. Modulating the association kinetics in this way, the SARS-COV-2 protein segment potentially encourages association of the monomers into a narrow distribution of only few oligomer types which may favor seeding of rod-like fibrils. However, SARS-COV-2 proteins can also enhance amyloid formation by a second mechanism. By raising the stability of the fibril polymorphs differentially, the Le Chatelier principle implies that interactions with viral proteins may shift the equilibrium between monomers, oligomers and fibrils toward specific fibril polymorphs. For this reason, we have performed a second set of molecular dynamics simulations studying the effect of SK9 on the stability of the two most common α-synuclein fibril structures^16^, the rod and the twister polymorphs of **Figure 1**. Our goal is to investigate whether the presence of SK9 stabilizes α-synuclein fibrils; and if yes, how this effect depends on the fibril geometry. As for the monomer simulations we compare for the two fibril polymorphs trajectories where SK9 fragments bind initially to the α-synuclein (but can change position or even detach), with our control, the corresponding simulations where the SK9 fragments are absent. Since only parts of the chain conformations have been resolved in the experimental models, and these models differ in length for rod and twister, we have also studied corresponding fibril models where missing residues are added to achieve equal chain length. Note, however, that in order to reduce the computational costs these models are not made of full-size α-synuclein chains but only of fragments 38-120. For details, see the method section.

A visual inspection of the final conformations in **Figure 6a-6d**, where for a more easy comparison we display only the residues 38-97, shows only marginal stabilization by SK9 for the rod fibril architecture when only the experimentally resolved fragment of residues 38 to 97 is simulated, and even some destabilization for the extended rod fibril (residues 38-120). The visual inspection is quantified by the time evolution in RMSD in **Figure 6e** and the residue-wise RMSF in **Figure 6f**. Both quantities are again calculated only for residues 38 to 97, i.e., ignoring the disordered and unresolved parts of the extended fibril model. Note, however, that the error bars are in both cases large, suggesting that the presence of SK9 has only little effect on the stability of the rod-fibril architecture. This is consistent with the rather small changes in SASA, packing distance between the folds, number of both the packing contacts (between folds) and stacking contacts (between the layers in each fold), and the frequencies of native inter- and intra-chain contacts, see **Table 2**.

**Table 2:** Mean values of total SASA, packing distance, number of intrachain, stacking and packing contacts, and frequency of native interchain and intrachain contacts. All values are averages over the final 50 ns of all trajectories of the respective systems. Note that we consider only for the experimentally resolved regions in our calculation of these quantities, i.e., ignoring the disordered and unresolved parts of the fibril models.

| system | | SASA ($\text{\AA}^2$) | packing distance ($\text{\AA}$) | intrachain contacts | stacking contacts | packing contacts | % of native interchain contacts | % of native intrachain contacts |
| --- | --- | --- | --- | --- | --- | --- | --- | --- |
| Rod fibril model (PDB-ID: 6CU7; residues 38-97) | control | 29472 (878) | 10.4 (0.7) | 56 (4) | 359 (4) | 27 (2) | 84 (6) | 60 (5) |
|  | with SK9 | 30490 (656) | 10.6 (0.7) | 53 (5) | 354 (5) | 23 (2) | 84 (6) | 60 (5) |
| Extended rod fibril (residues 38-120) | control | 22514 (423) | 10.0 (0.9) | 56 (3) | 350 (17) | 25 (3) | 82 (4) | 60 (4) |
|  | with SK9 | 27874 (685) | 10.6 (0.6) | 62 (7) | 332 (6) | 24 (2) | 77 (6) | 60 (4) |
| Twister fibril model PDB-ID: 6CU8; residues 43-83) | control | 20918 (615) | 10.6 (0.4) | 19 (2) | 221 (6) | 42 (2) | 80 (6) | 46 (4) |
|  | with SK9 | 21717 (485) | 10.3 (0.7) | 17 (3) | 220 (3) | 40 (1) | 81 (4) | 46 (4) |
| Extended twister fibril model (residues 38-120) | control | 15008 (389) | 9.6 (0.8) | 21 (3) | 215 (5) | 38 (2) | 80 (6) | 44 (5) |
|  | with SK9 | 19882 (771) | 9.8 (0.8) | 13 (6) | 213 (10) | 38 (3) | 80 (6) | 35 (5) |

On the other hand, both visual inspection and analysis of RMSD and RMSF in the corresponding **Figure 7** show a signal for stabilization of the twister α-synuclein fibril structure in the simulations that started with SK9 bound to the fibrils, but again no clear signal is seen for this stabilization in the quantities listed in **Table 2**. This may be because the effect of SK9 is localized: the RMSF plots in **Figure 7f** suggests a lower flexibility of the region 45-65 in simulations where the viral protein fragment SK9 is present. This region is one of the main binding-partners for SK9, see **Figure 8** where we show the binding probability as measured in simulations starting from fibril models with non-extended chains (the corresponding figures for the extended fibril models are shown as **Supplemental Figure S3**).

**Figure 7:**
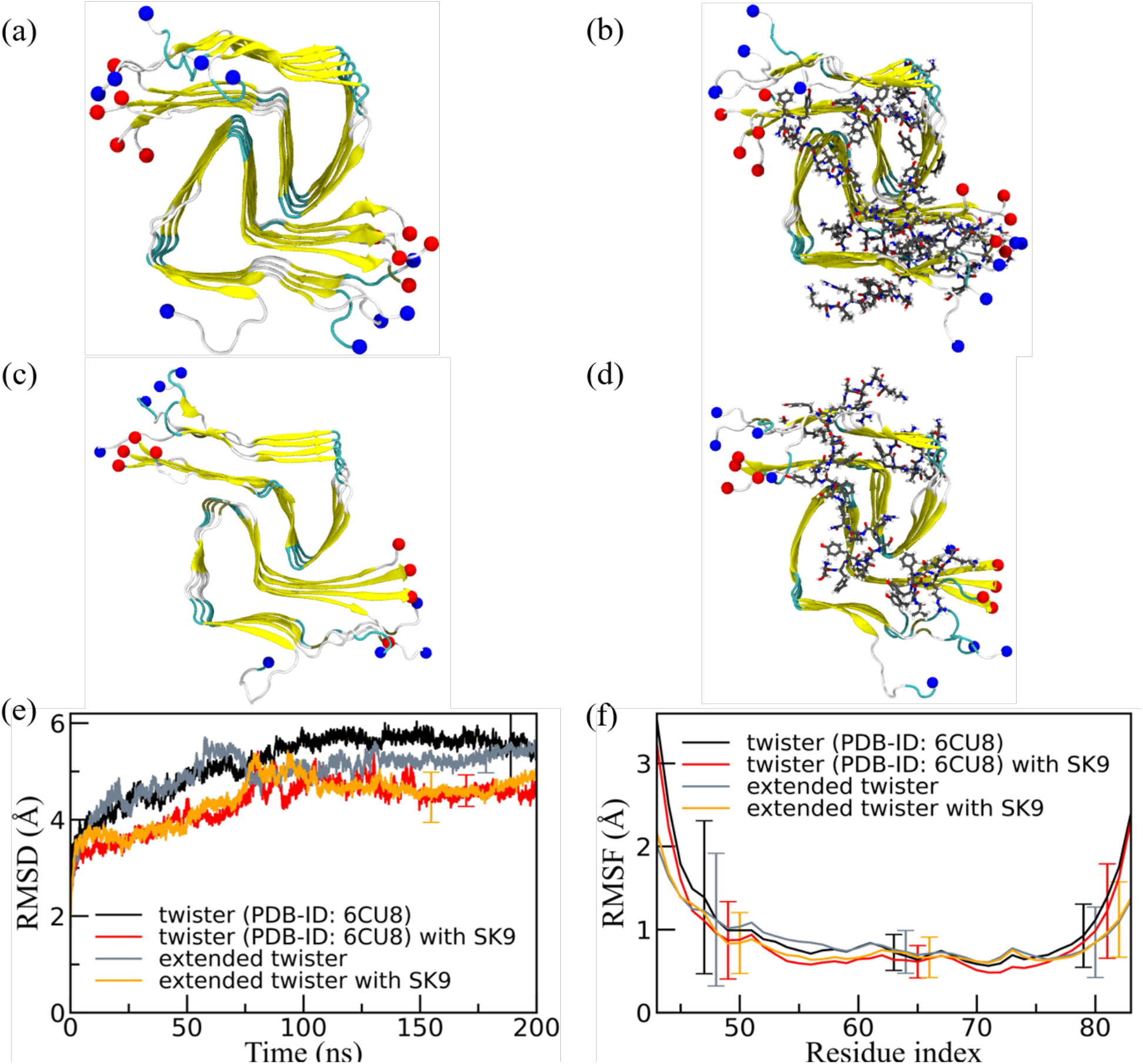
Representative final configurations extracted from simulations starting from (a) the experimentally determined twister-like α-synuclein fibril model (PDB-ID: 6CU8) and (c) the extended model. Corresponding final snapshots extracted from simulations in the presence of SK9-segment are shown in (b), and (d). N- and C-terminus are represented by blue and red spheres, respectively. Only residues 43-83 are shown for the extended model configurations in (c) and(d). The time evolution of the RMSD in the simulation of these systems is shown in (e), and residue-wise RMSF in (f). We calculate RMSD and RMSF again only for the experimentally resolved region 43-83, i.e., ignoring the disordered and unresolved parts of the fibril models, considering all backbone atoms. Only a few typical errorbars are shown to make figures more readable.

**Figure 8:**
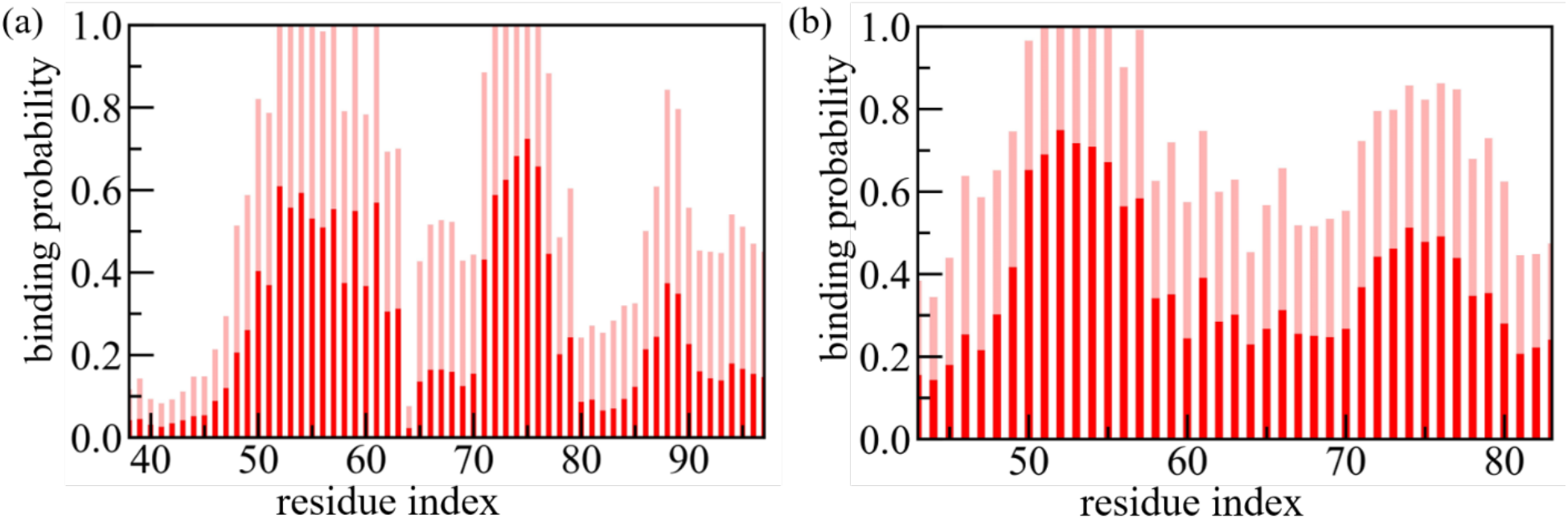
The residue-wise normalized binding probability of the SK9 segment for α-synuclein rod (a) and twister (b) polymorph. Data are averaged over the final 50 ns of each trajectory and shaded region represents the standard deviation.

In order to understand in more detail changes in fibril stability upon addition of SK9, we have looked into the change of contact pattern in both fibril geometries. In **Figure 9** we show contact probabilities for stacking contacts (i.e., such between residues located on neighboring layers of the same fold) for the rod-like fibril in the absence (**Figure 9a**) and in the presence of (**Figure 9b**) the SK9 segment. The corresponding probabilities for the twister fibril are shown in **Figure 9d** (absence of SK9) and in **Figure 9e** (presence of the SK9 segment). The differences between probabilities measured in the presence and absence of SK9 are shown in **Figures 9c** (rod fibril) and **Figure 9f** (twister fibril). The data shown in **Figure 9** are from the simulations using the same chain-length as in the experimentally determined fibril models. The corresponding probabilities as measured in simulations of the fibrils with extended chains are shown in **Supplemental Figure S4**. As shown already in **Table 2**, native interchain contacts are more preserved than intrachain contacts. Most of the native contact pairs in the NAC region are highly conserved over the simulation, but hydrophobic contacts involving F94, the T59-T72 contacts and such with the G67 and G68 residues in the turn region are seen with a frequency of less than 10%. However, with the exception of a gain of contacts between residues in the segment of G68-A78 with the segment of residues I88-K97 no signal for changes in the stacking pattern upon binding of SK9 is found for the rod-like fibril. The exception can be explained by the preferential binding of SK9 to residues V70-A78, which stabilized this segment (see also the difference in intrachain contacts in **Figure 10c**) and leads to the observed stacking contacts.

**Figure 9:**
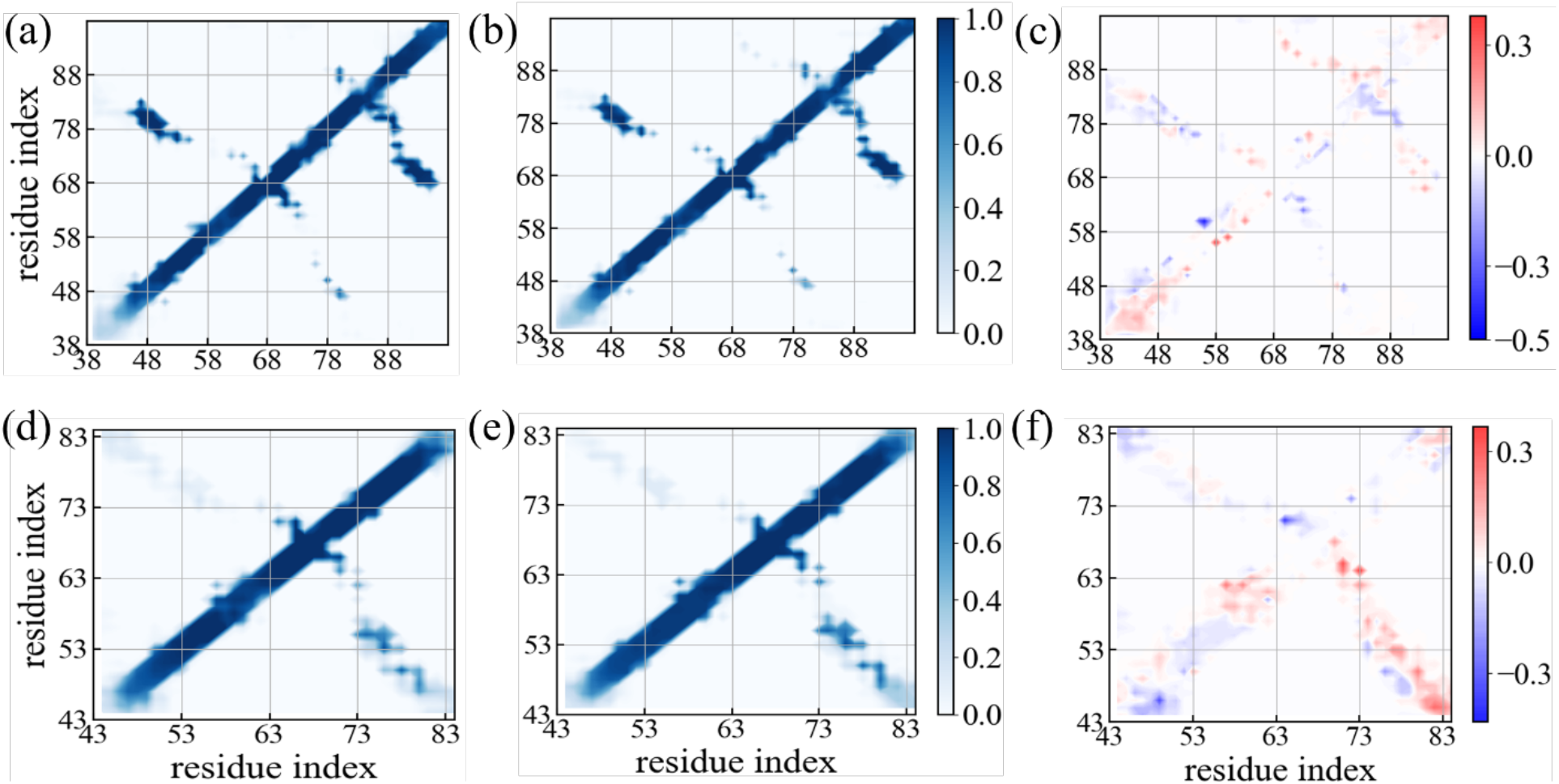
Residue-wise stacking contact probabilities measured in simulations of the rod-like fibril in the absence (a) and in the presence of the SK9 segment (b). Corresponding data measured in simulations of the twister-like fibril are shown in (d) and (e). The differences in contact probability resulting from the presence of SK9 are shown in (c) and (f), respectively. Data are averaged over the final 50 ns of each trajectory.

**Figure 10:**
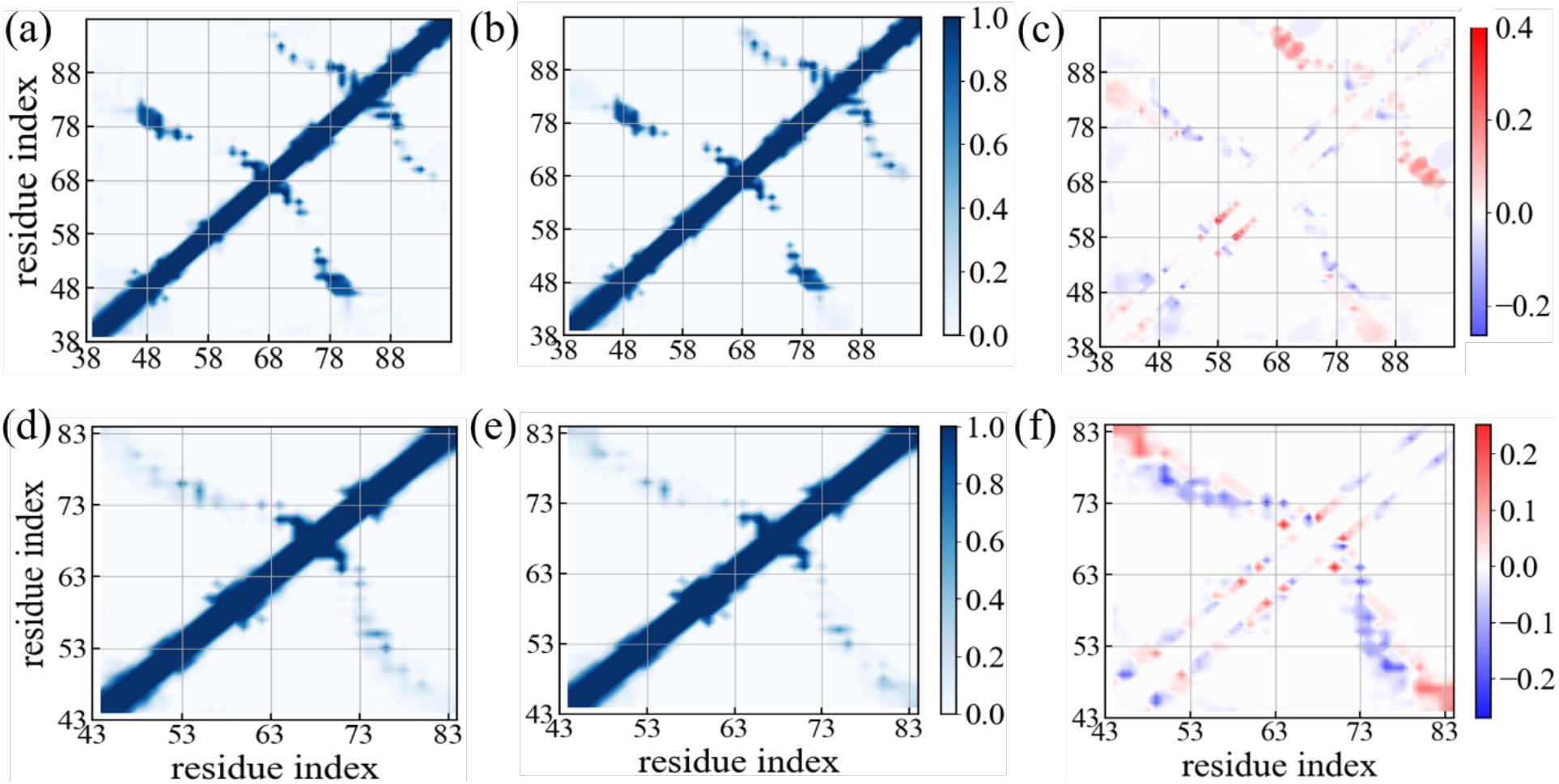
Residue-wise intrachain contact probabilities measured in simulations of the rod-like fibril in the absence (a) and in the presence of the SK9 segment (b). Corresponding data measured in simulations of the twister-like fibril are shown in (d) and (e). The differences in contact probability resulting from the presence of SK9 are shown in (c) and (f), respectively. Data are averaged over the final 50 ns of each trajectory.

On the other hand, while native interchain contacts are only slightly less stable in the twister fibril than in the rod, we find a larger reduction in the frequency of native intrachains, see **Table 2**. Especially, native hydrophobic contacts involving the residue pairs V66-A69, V66-A70 and V74-V77, and native contacts involving glycine residues (G67, G73) in the inter-protofilament interface are disrupted. Upon SK9-binding, we observe an significant increase in stacking contacts between residues of the segment K45-N65 with that of the segment G73-K80, which is compensated by a pronounced loss in the intrachain contacts in the same region, see **Figure 10** and **Supplemental Figure S5**. Hence, interaction with SK9 leads in the Twister fibril to a replacement of contacts between residues within chains by contacts between residues located on chains in neighboring layers. This may explain why the observed stabilization of the twister fibril geometry upon binding of SK9, seen in the RMSD plots, does not lead to more pronounced changes in the quantities shown in **Table 2**.

## Conclusions

As virus-induced amyloid-formation has been observed *in vitro*^5^, and may suggest onset of Parkinson’s Disease and related neurodegenerative diseases^2–5^ as potential long-term risk for COVID-19 patients, it is important to investigate how amyloidogenic regions of SARS-COV-2 initiate or accelerate formation of α-synuclein aggregates, the cytotoxic agent in Parkinson’s Disease.^6^ For this purpose, we have studied, using all-atom molecular dynamics simulations, how interaction with a short segment (SK9) of the Envelope protein of SARS-COV-2 changes the conformational ensemble of α-synuclein chains and the stability of two α-synuclein fibril polymorphs, the rod and the twister structure.

We find that the presence of SK9 changes the ensemble of α-synuclein not only toward more aggregation-prone conformations but that this effect likely leads to a preference for the rod fibril motif. Crucial for this effect is a change in the contact pattern that causes higher flexibility, reduced helix-propensity and larger exposure of residues, especially in the segment E46-A56 that form in the rod fibril polymorph (see **Figure 6**) the inter-protofilament interface. Especially important is the increase infrequency and lifetime of the two contacts E46-K80 and V52-A76. Both are found also in the experimentally resolved rod-fibril form, and the mutation E46K, which destroys the salt-bridge E46-K80, is one of the familial mutations that cause Parkinson’s Disease and that shift the equilibrium from rod to twister fibrils.^16^

The outcome of our fibril simulations is less obvious. While we see some stabilization of the fibril geometries in the presence of SK9, the signal is weak, and it is only for the twister form correlated with a clear change in the contact pattern. Hence, it appears that the viral protein fragment SK9 may alter α-synuclein amyloid formation but has little effect of the stability of pre-existing or newly-formed fibrils.

The preference for monomer conformations that likely seed the rod-like fibrils is interesting as the twister fibril is considered to be more cytotoxic.^16^ Further simulations, using other amyloidogenic segments of SARS-COV-2 proteins, especially in the Spike-protein, are needed to see if this effect is specific for the SK9 segment. It will be especially interesting to compare our work with recent studies into the amyloidogenesis of Spike-protein fragments after endoproteolysis.^39^

## AUTHOR INFORMATION

### Notes

The authors declare no competing financial interest.

## Acknowledgment

The simulations in this work were done using the SCHOONER cluster of the University of Oklahoma, XSEDE resources allocated under grant MCB160005 (National Science Foundation), and TACC resources allocated under grant under grant MCB20016 (National Science Foundation). We acknowledge financial support from the National Institutes of Health under grant GM120634.

