## Supplemental Figures for "Effect of an amyloidogenic SARS-COV-2 protein fragment on α-synuclein monomers and fibrils"

### Supporting Information

Supporting information contains five supporting figures (S1-S5).

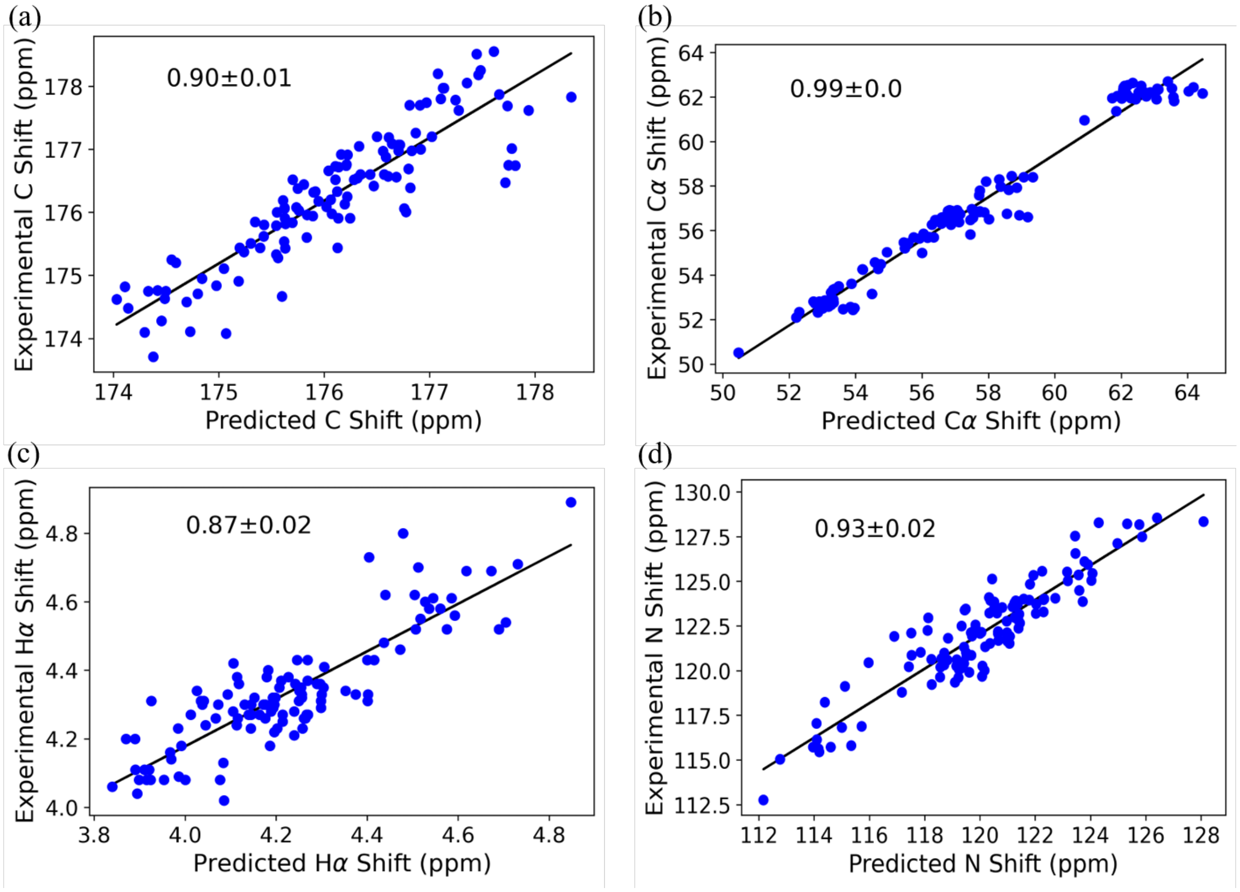


**Figure S1:** Regression plots of experimental (BMRB database entry 6968) over calculated C, Cα N and Hα chemical shifts for wild-type full-length α-synuclein monomer simulations are shown in (a)-(d). The Pearson correlation coefficients are also shown.

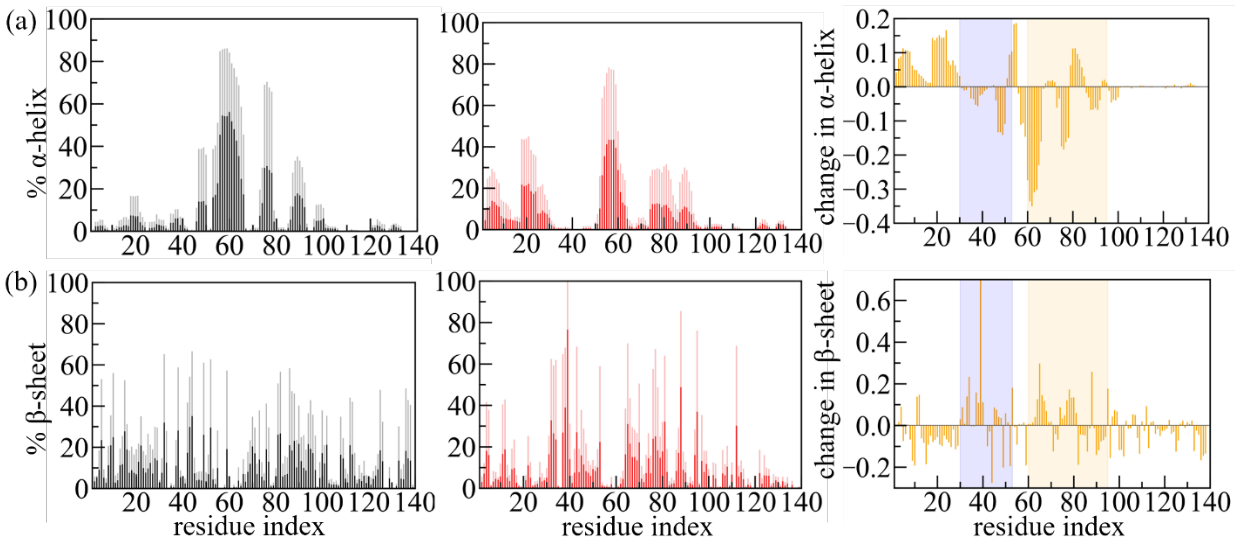


**Figure S2:** Residue-based α-helix (a) and β-sheet (b) propensity of wild-type α-synuclein monomer in the absence (black, left panel)) and in the presence (red, central panel)) of the SK9-segment. The difference in the respective residue-wise propensities is drawn in the right panel (yellow). Familial mutation sites (A30-A53) and NAC regions (K60-V95) are highlighted in light blue and yellow, respectively.

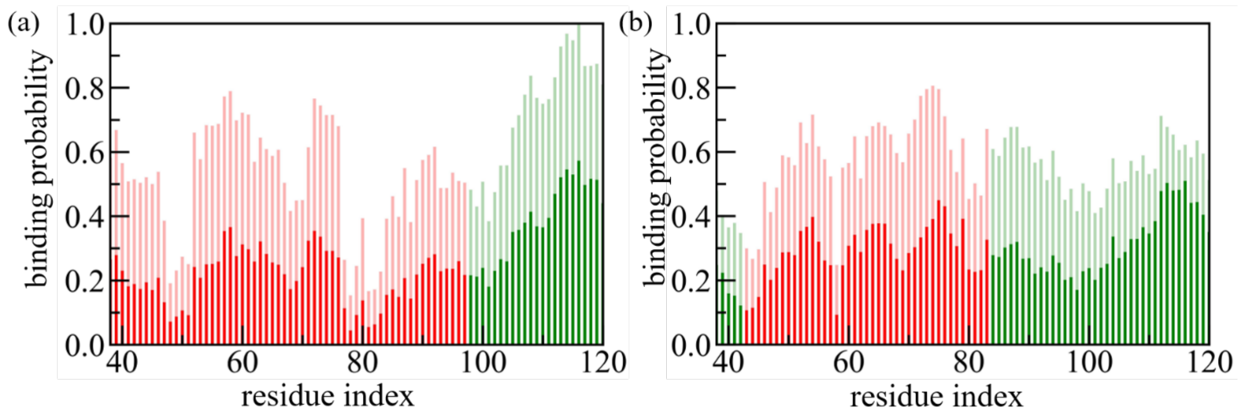


**Figure S3:** The residue-wise normalized binding probability of the SK9 segment for α-synuclein rod (a) and twister (b) polymorphs, where the individual chains are extended to residues 38-120. Data are averaged over the final 50 ns of each trajectory and shaded region represents the standard deviation. Binding frequencies for the experimentally resolved segments are colored in red, while binding frequencies for the unresolved parts are drawn in green.

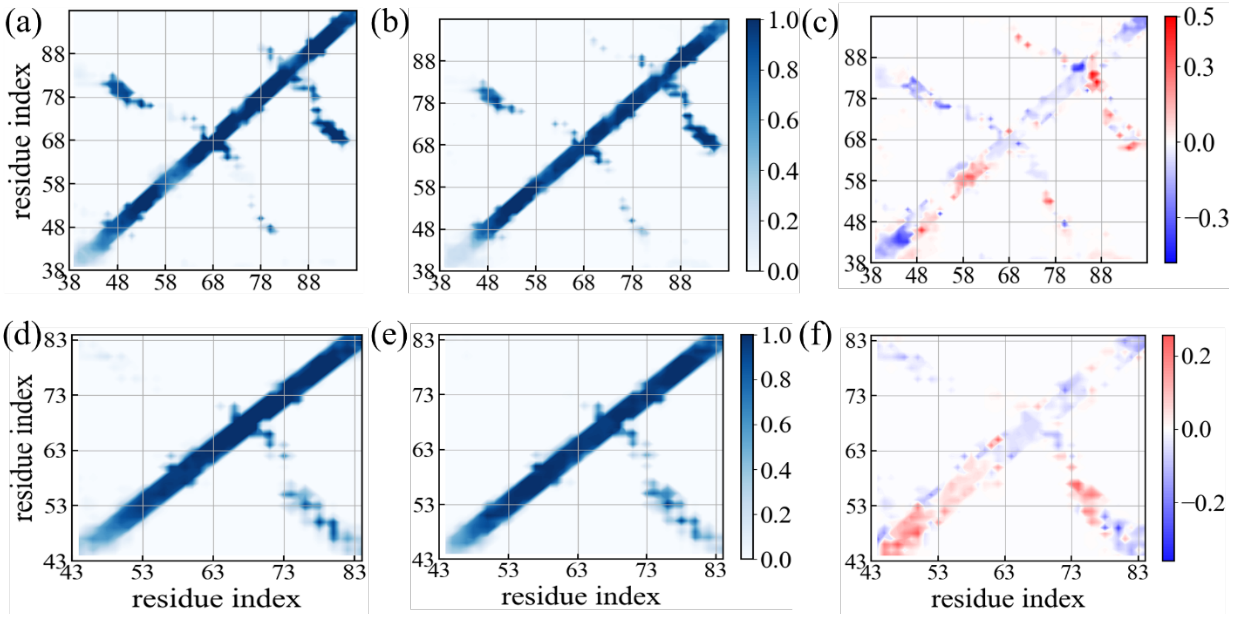


**Figure S4:** Residue-wise stacking contact probabilities measured in simulations of the rod-like fibril in the absence (a) and in the presence (b) of the SK9 segment. Corresponding data measured in simulations of the twister-like fibril are shown in (d) and (e). The differences in contact probability resulting from the presence of SK9 are shown in (c) and (f), respectively. Data are from simulations where the individual chains are extended to residues 38-120. Data are averages over the final 50 ns of each trajectory, and are calculated only for the experimentally resolved regions.

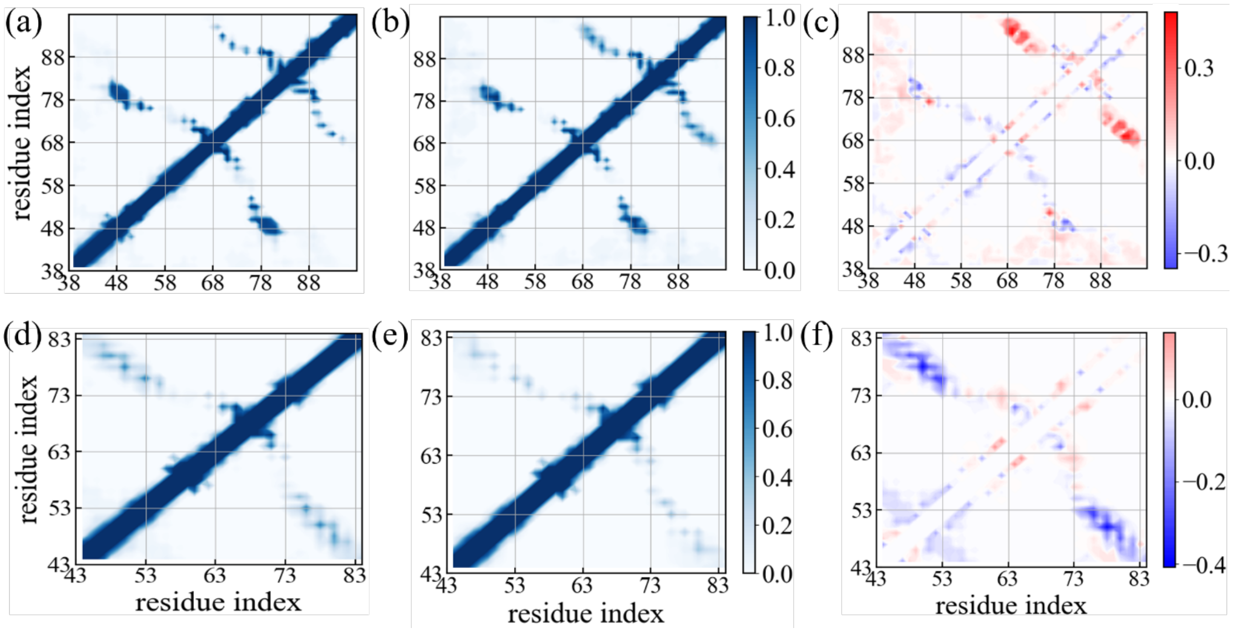


**Figure S5:** Residue-wise intrachain contact probabilities measured in simulations of the rod-like fibril in the absence (a) and in the presence (b) of the SK9 segment. Corresponding data measured in simulations of the twister-like fibril are shown in (d) and (e). The differences in contact probability resulting from the presence of SK9 are shown in (c) and (f), respectively. Data are from simulations where the individual chains are extended to residues 38-120.Data are averages over the final 50 ns of each trajectory, and are calculated only for the experimentally resolved regions.
